# Recognition of sounds by ensembles of proteinoids

**DOI:** 10.1101/2023.07.17.549338

**Authors:** Panagiotis Mougkogiannis, Andrew Adamatzky

## Abstract

Proteinoids are artificial polymers that imitate certain characteristics of natural proteins, including self-organization, catalytic activity, and responsiveness to external stimuli. This paper investigates the potential of proteinoids as organic audio signal processors. We convert sounds of English alphabet into waveforms of electrical potential, feed the waveforms into proteinoid solutions and record electrical responses of the proteinoids. We also undertake a detailed comparison of proteinoids’ electrical responses (frequencies, periods, and amplitudes) with original input signals. We found that responses of proteinoids are less regular, lower dominant frequency, wider distribution of proteinoids and less skewed distribution of amplitudes compared with input signals. We found that letter of English alphabet uniquely maps onto a pattern of electrical activity of a proteinoid ensemble, that is the proteinoid ensembles recognise spoken letters of English alphabet. The finding will be used in further designs of organic electronic devices, based on ensembles of proteinoids, for sound processing and speech recognition.

## 1. Introduction

Proteinoids are protein-like molecules that can be synthesised abiotically from amino acids in specific environments, such as high temperature or low pH. These molecules have been suggested as potential precursors to the earliest living cells, as they can self-assemble into microspheres that display certain life-like properties, including osmosis, budding, and fission [1, 2, 3, 4, 5, 6, 7, 8, 9, 10]. This paper investigates the potential use of proteinoids as bioinspired audio signal processors. Our hypothesis suggests that proteinoids possess the ability to function as nonlinear filters, enabling them to modify and enhance audio signals through processes such as amplification, distortion, compression, and modulation [11, 12]. Our hypothesis is derived from the analogy between proteinoids and biological membranes, both of which exhibit electrical properties and respond to acoustic stimuli [13, 14]. In order to evaluate our hypothesis, we synthesised diverse proteinoids using various amino acid mixtures and subjected them to varying audio signals. Subsequently, we conducted measurements and performed analysis on the output signals utilising diverse techniques and metrics.

The paper is structured as follows: Section 2 outlines the methods and materials employed in the synthesis and characterization of proteinoids. Section 3 presents the results and analysis of the audio signal processing experiments. Section 4 explores the implications and applications of our findings. Finally, Section 5 concludes the paper and proposes potential avenues for future research.

## 2. Methods

L-Phenylalanine, L-Aspartic acid, L-Histidine, L-Glutamic acid, and L-Lysine were obtained from Sigma Aldrich with a purity exceeding 98%. Proteinoids were synthesised using established techniques described in a previous study [15] (Fig. 1). The proteinoids’ structures were examined using FEI Quanta 650 equipment for scanning electron microscopy (SEM). The proteinoids were characterised using FT-IR spectroscopy [15]. The synthesis of proteinoids L-Glu:L-Arg from the amino acids glutamic acid and arginine was carried out according to the procedure outlined in Fig. 2. The proteinoid chains were obtained by heating stirred the mixture of amino acids at a temperature of 200°C for a duration of 15 minutes. The proteinoid microspheres were formed by adding water and stirring the proteinoid chains for an additional 15 minutes. The proteinoid microspheres were filtered, washed, and subsequently freeze-dried to obtain a dry sample of proteinoids L-Glu:L-Arg.

**Figure 1:**
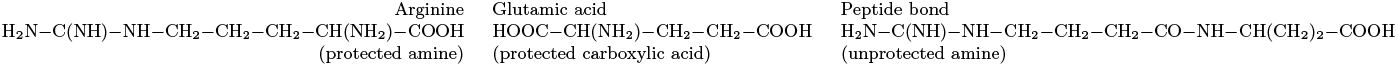
The process of coupling two amino acids in a solution. The amine group of one molecule reacts with the carboxylic acid group of the other molecule to form a peptide bond.

**Figure 2:**
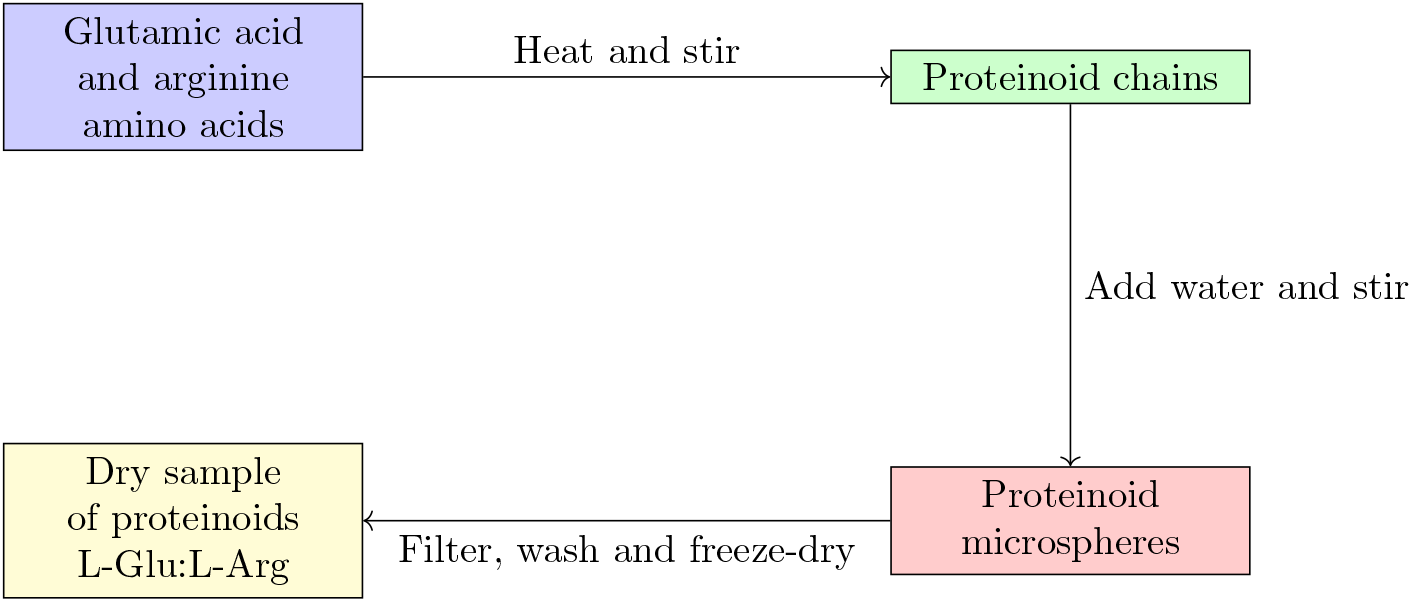
Here is a schematic diagram illustrating the synthesis of proteinoids L-Glu:L-Arg from the amino acids glutamic acid and arginine. The process includes heating and stirring the mixture of amino acids, followed by the addition of water and further stirring to create proteinoid microspheres. The next steps involve filtering, washing, and freeze-drying the sample to obtain the proteinoids in a dry form.

A high-resolution data logger with a 24-bit A/D converter (ADC-24, Pico Technology, UK) and iridium-coated stainless steel sub-dermal needle electrodes (Spes Medica S.r.l., Italy) were used to measure the electrical activity of the proteinoids. To assess the potential difference, pairs of electrodes were set up with a spacing of about 10 mm between them. At a rate of one sample per second, all electrical activity was captured. Multiple measurements (up to 600 per second) were captured by the data logger, and their average was saved for further study.

The BK 4060B function generator, further in the paper referred to as ‘device’, was utilised for the generation of electrical spikes. The device is a function/arbitrary waveform generator with dual channels. It has the capability to accurately generate various waveforms such as sine, square, triangle, pulse, and arbitrary waveforms. The maximum waveform generation rate is 75 MSa/s in real point-by-point arbitrary mode and 300 MSa/s in direct digital synthesis mode.

An experimental setup was designed to analyse the response of proteinoids to audio signals, as depicted in Fig. 3. The laptop’s microphone was used to capture the audio signals, which were then processed by Matlab. The result of this processing was CSV files that contained a single column of potential values. The audio signals included the letters A to Z of the English alphabet. These letters were pronounced by a male speaker using the vocal database found at https://freesound.org/people/dersuperanton/sounds/434730/. This database includes recordings of individual letters spoken both in isolation and with various tones. Next, we utilised electrodes connected to a BK precision 4053 B transfer function to apply the CSV audio files to the proteinoids. The response of proteinoids to audio signals was recorded using a Picoscope 4000 series oscilloscope. The resulting data was saved as CSV files for subsequent analysis.

**Figure 3:**
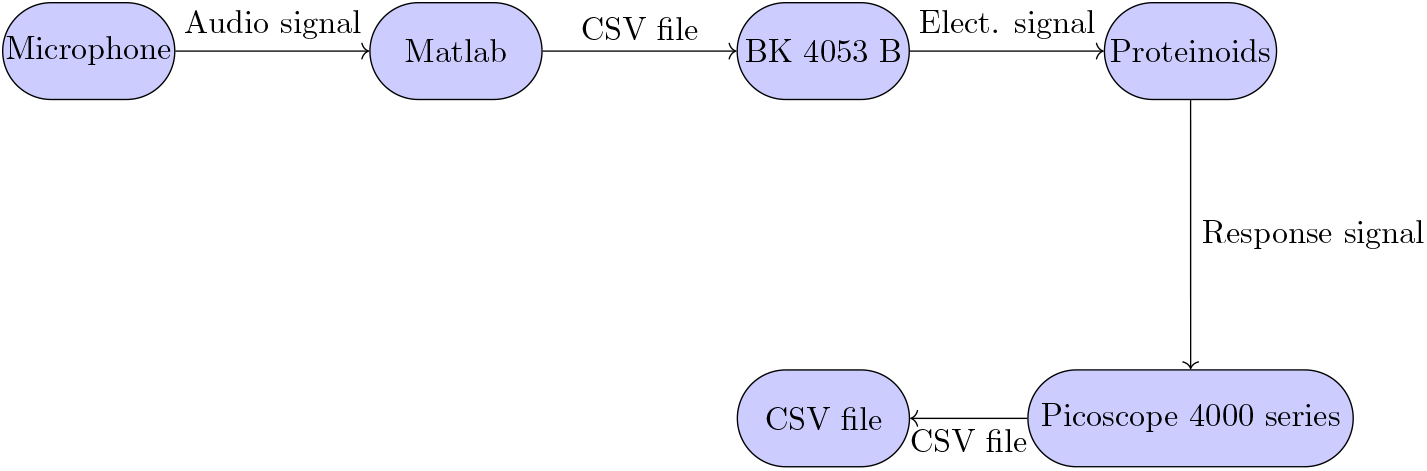
Schematic diagram of the experimental setup for analyzing the response of proteinoids to audio signals.

We utilised Matlab to develop a script that effectively simulates the spherical compression wave of the proteinoids. This script calculates and plots the pressure, taking into account the variables of radius and time. The script utilises the dominant frequency and maximum potential of the proteinoid signal as parameters for the sound wave, along with the speed of sound in air. The script assumes that the sound waves start with a phase of zero and propagate in a radial manner from the source. The script generates two surface plots: one for the device signal and another for the proteinoid signal. These plots display the real part of the pressure at various distances and times.

The microphone recorded the acoustic signals of the proteinoid and the device, which were then converted into potential vectors. The potential vectors were quantized into 4-bit digital signals using 16 levels and a range of [-1, 1]. The discrete values of the digital signals were plotted as stem plots with markers, illustrating their variation over time. The digital signals capture both the amplitude and frequency modulation of the acoustic signals, along with any noise and variations present in the recording. Figure 4 presents the diagram illustrating the digital signals.

**Figure 4:**
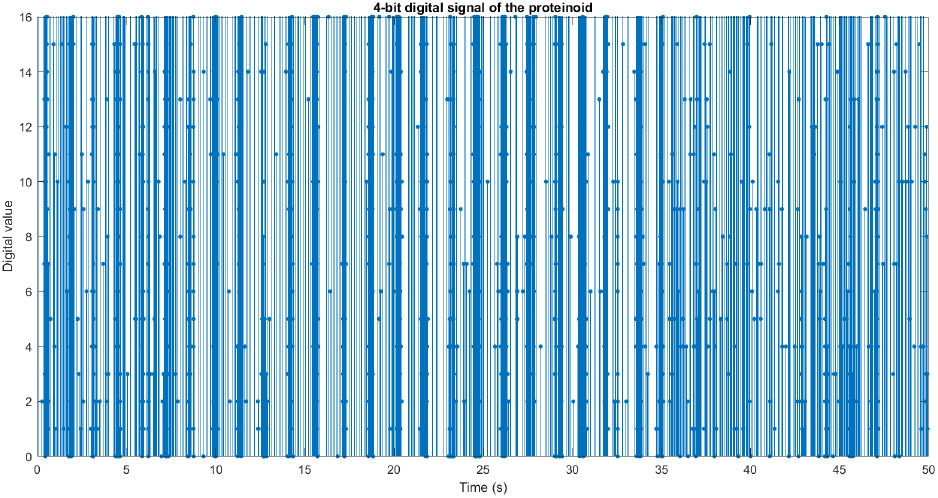
The proteinoid’s graphical representation of a 4-bit digital sound wave. A microphone captured the sound wave and converted it into a potential vector. The potential vector was then quantized to a 4-bit digital signal with 16 levels and a range of [-1 1] to create a 4-bit digital signal. The discrete signal values over time were displayed as a stem plot with markers for the digital signal. The digital signal reflects the amplitude and frequency modulation of the sound wave, along with the recording’s noise and variation.

## 3. Results

The morphology of proteinoid ensembles was examined using scanning electron microscopy (SEM). Figure 5 shows a scanning electron microscope (SEM) image of proteinoid nanoparticles, exhibiting a diameter of approximately 20 nanometers and possessing a uniformly smooth surface. The image was captured at a magnification of 60,000x and an applied voltage of 2 kV. In their early stages, these microspheres display a spherical shape and have a smooth surface. These findings are consistent with previous studies on proteinoid microspheres and their potential importance in the origin of life [5, 16]. The image suggests that proteinoid microspheres have diverse surface structures and textures, which may impact their conductivity and habituation properties as neuromorphic computing devices [17].

**Figure 5:**
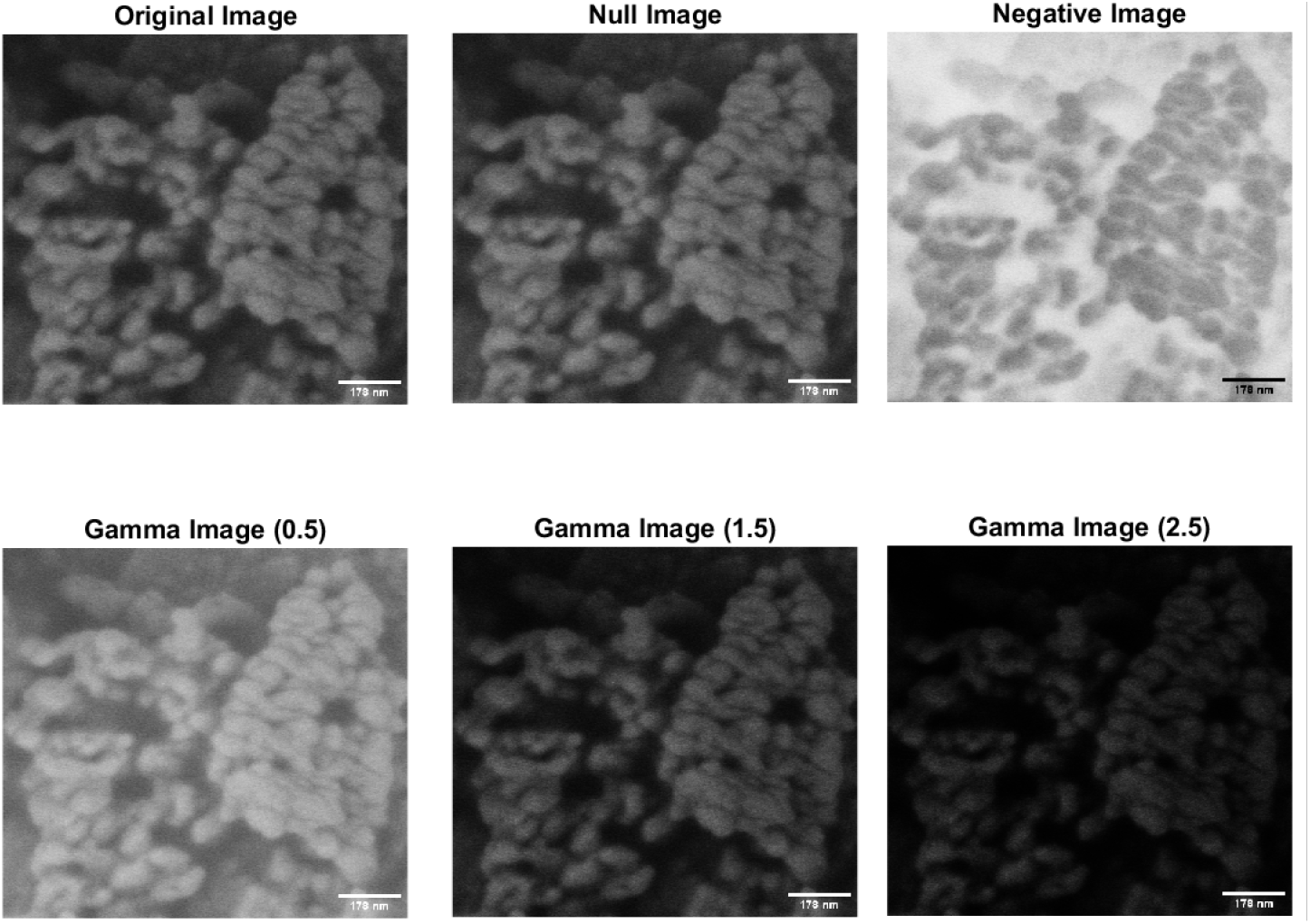
The provided image depicts proteinoid nanoparticles obtained through the process of heating a mixture of L-glutamic acid and L-arginine to their boiling point, as observed through scanning electron microscopy (SEM). The nanoparticles have a diameter of around 20 nanometers and a smooth surface. The image was obtained at a magnification of 60,000x and a voltage of 2 kV was applied. Proteinoid microspheres are being considered as a viable model for studying the origins of life and potential applications in neuromorphic computing. The image was subjected to different transformation techniques in order to enhance contrast and improve visibility. The null image represents the unmodified source image. The negative image is created by inverting the original image, which involves subtracting the intensity values from 255. Gamma images are generated by applying a non-linear function with different gamma values to the original images. A gamma value less than 1 increases image brightness, while a gamma value greater than 1 decreases image brightness. The gamma images were created in MATLAB using the imadjust function with gamma values of 0.5, 1.5, and 2.5.

As depicted in Fig. 6, the potentials of the device and the proteinoid differ in response to an audio signal containing the English alphabet. The device potential varies with the frequency and amplitude of the audio signal, whereas the proteinoid potential displays a distinct pattern. This indicates that proteinoid provide distinctive responses to audio signals converted to electrical waveforms.

**Figure 6:**
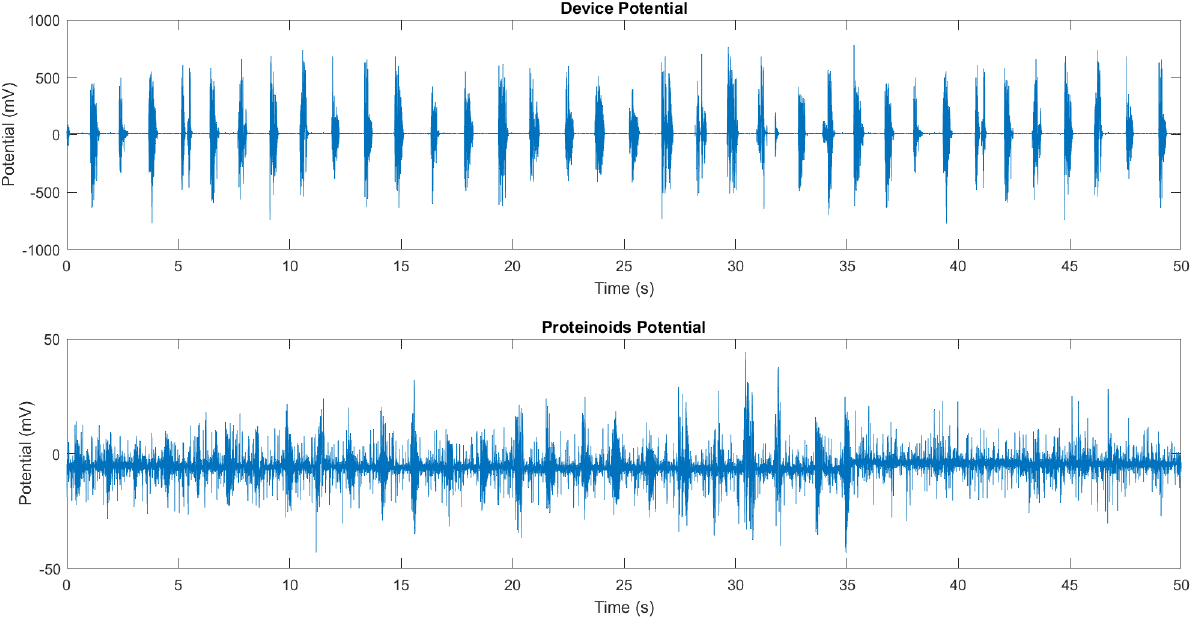
The responsiveness of a device BK PRECISION 4053 B and a proteinoid to an acoustic signal containing the English alphabet. The graph depicts the waveform of the audio signal, which consists of 26 consecutively spoken letters. The potential of the proteinoid, displayed in millivolts (mV), also fluctuates in response to the audio input, albeit in a different manner (see bottom panel). The image depicts differences in response to identical stimuli between the device and the proteinoid.

Figure 7 shows that the linear relationship between the two signals is extremely weak, with a correlation coefficient of 7.0656e-04. The null hypothesis of no connection is rejected with a p-value of 0, indicating that there is a substantial deviation from zero. However, this does not prove that one signal causes the other. Comparing the device’s average voltage of -5.3161 mV to the proteinoid’s average voltage of 10.8667 mV reveals that the device has a lower average voltage.

**Figure 7:**
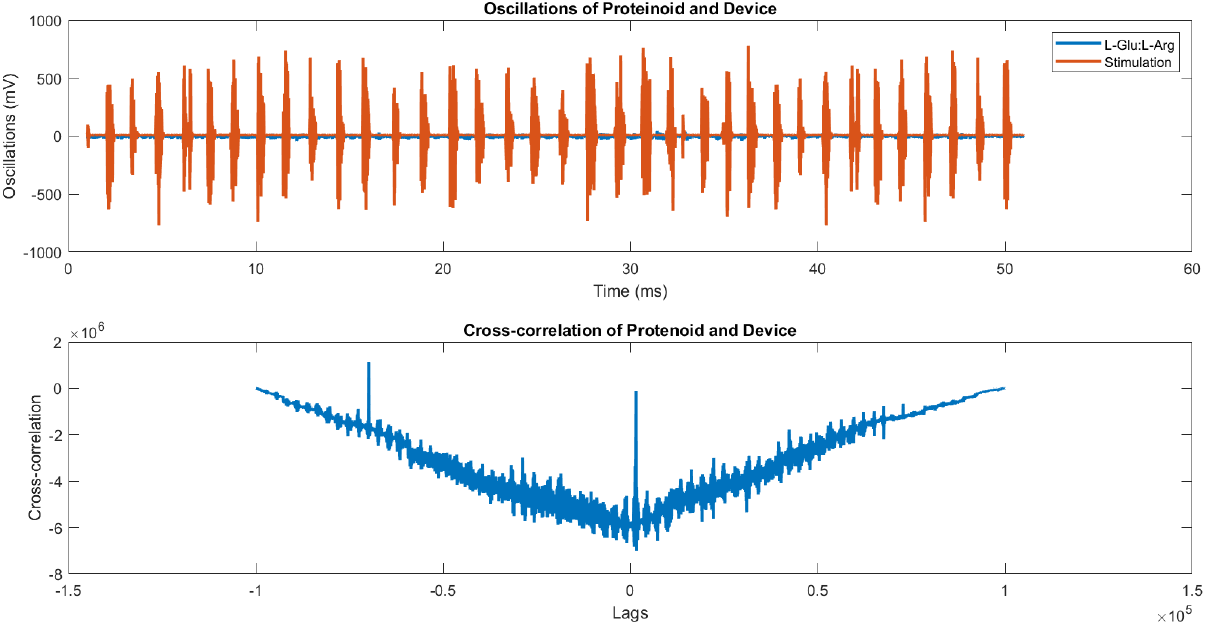
Proteinoid and device oscillations, as well as their association with one another. Time series of proteinoid (blue) and device (red) voltage readings are displayed above. In the bottom panel, we can see the cross-correlation function between the two signals.

The analysis aimed to determine the statistical significance of the mean difference between two groups, device and proteinoid, as presented in Table 1. The F-statistic is computed as the ratio of the between-group mean square (MS) to the within-group mean square (MS). It quantifies the degree of heterogeneity among group means in relation to the heterogeneity within each group. A high F-statistic suggests significant differences between group means relative to within-group variability. The p-value represents the likelihood of obtaining an F-statistic that is equal to or greater than the observed value, assuming the null hypothesis that the means of the groups are equivalent. A small p-value suggests rejection of the null hypothesis and supports the presence of a significant difference between group means. The F-statistic and p-value in this instance are highly significant, with values of 2.4661e+03 and 0, respectively. The null hypothesis is rejected, indicating a significant difference between the means of device and protenoids.

**Table 1:** An ANOVA table was constructed to compare the means of two groups, namely device and proteinoid. The statistical analysis shows a significant difference between the group means, as evidenced by the F-statistic and p-value.

| Source | SS | df | MS | F | Prob>F |
| --- | --- | --- | --- | --- | --- |
| Columns | 1.3095e+07 | 1 | 1.3095e+07 | 2.4661e+03 | 0 |
| Error | 1.0620e+09 | 200006 | 5.3099e+03 | - | - |
| Total | 1.0751e+09 | 200007 | - | - | - |

The temporal evolution of the device and proteinoid potentials was examined through the utilisation of histograms and Fourier transforms (Figure 8 and Figure 9). The device potential exhibited a distinct periodic pattern characterised by regular peaks exceeding 20 mV. In contrast, the proteinoid potential displayed a more irregular pattern with fewer peaks surpassing 20 mV. The Fourier transform of the device potential indicated a dominant frequency of 20.48 Hz, suggesting a high oscillation rate. The proteinoid potential’s Fourier transform showed a dominant frequency of 6.48 Hz, suggesting a lower oscillation rate. The histograms of the periods and amplitudes of the device and proteinoid potentials exhibited distinct distributions. The device potential exhibited a relatively narrow distribution of periods, with a mean of 2804.60 ms and a standard deviation of 314.12 ms, suggesting a reduced level of variability in the cycle length. The device potential exhibited a non-normal distribution of amplitudes, with a mean of 589.03 mV and a standard deviation of 131.58 mV, suggesting greater variability in the peak height. The proteinoid potential exhibited a broader range of periods, with an average of 2747.03 ms and a standard deviation of 546.42 ms, suggesting greater variability in the duration of the cycles. The proteinoid potential exhibited a more evenly distributed range of amplitudes, with an average of 18.23 mV and a standard deviation of 9.28 mV. This suggests a reduced level of variability in the height of the peaks.

**Figure 8:**
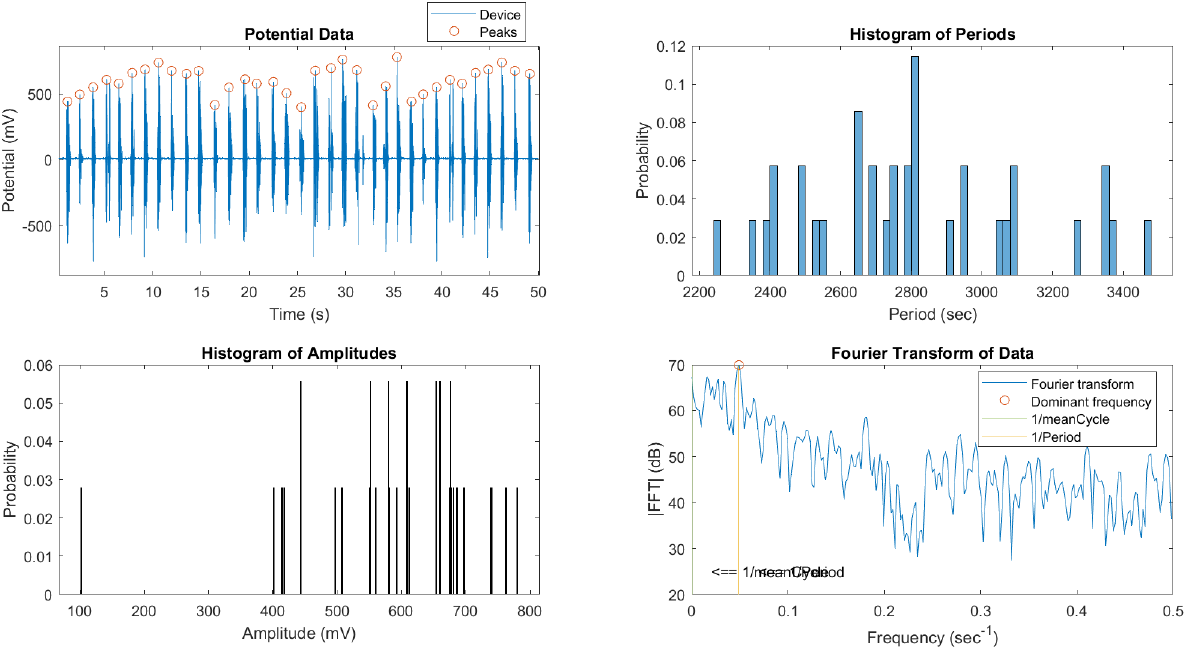
Data for the device’s potential, histogram of periods, histogram of amplitudes, and Fourier transform. The device potential exhibited a periodic pattern with peaks exceeding 20 mV and a frequency of 20.48 Hz. The period histogram displayed a limited distribution around the mean of 2804.60 milliseconds. The amplitude histogram displayed a biased distribution with a mean of 589.03 mV.

**Figure 9:**
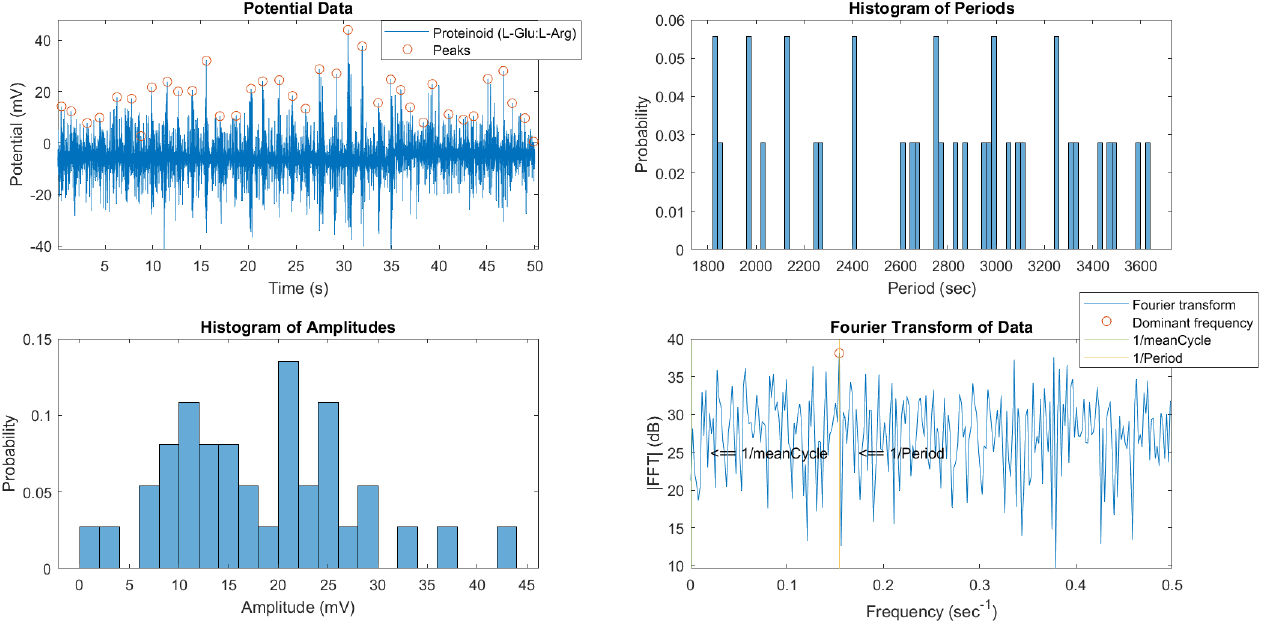
Data for the proteinoid’s potential, histogram of periods, histogram of amplitudes, and Fourier transform. Proteinoid potential exhibited a less periodic pattern with peaks exceeding 20 mV and a dominant frequency of 6.48 Hz. The period histogram displayed a wider distribution around the mean of 2747.03 milliseconds. The distribution of amplitudes in the histogram was more uniform, with a mean of 18.23 mV.

We compared the distributions of the peaks and periods of the device and proteinoid potentials by analysing their skewness and kurtosis values using box plots (Figure 10 and Figure 11). The device potential exhibited a negative skewness of -1.46 and a positive kurtosis of 6.35 for the peaks. In contrast, the proteinoid potential displayed a positive skewness of 0.54 and a positive kurtosis of 3.33 for the peaks. The device potential exhibited a mean skewness of 0.39 and a mean kurtosis of 2.47, whereas the proteinoid potential had a mean skewness of -0.20 and a mean kurtosis of 1.91. The results suggest that the device potential exhibited a left-skewed and peaked distribution of peaks, while the proteinoid potential showed a right-skewed and flat distribution of periods. The proteinoid potential exhibited a distribution of peaks that was more right-skewed and flat, and a distribution of periods that was more left-skewed and flat, compared to the device potential.

**Figure 10:**
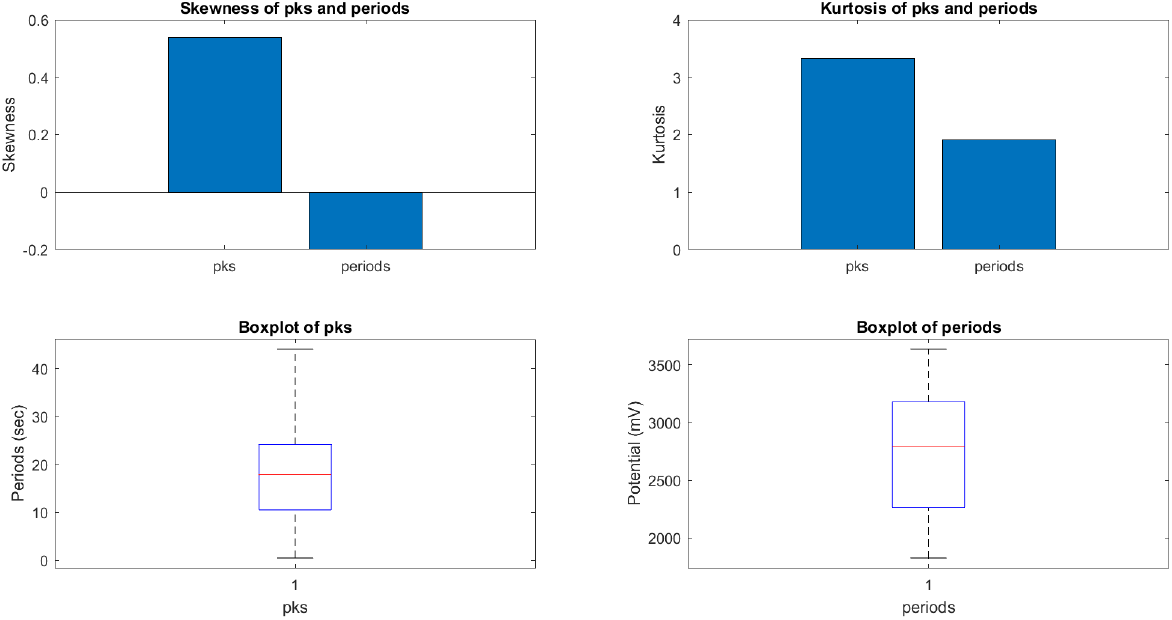
The box plots display the skewness and kurtosis of peaks and periods in the proteinoid potential. The proteinoid potential exhibited a positively skewed distribution with low kurtosis, indicating a right-skewed and platykurtic pattern for the peaks. The proteinoid potential exhibited a left-skewed and platykurtic distribution, as evidenced by its negative skewness and low kurtosis values.

**Figure 11:**
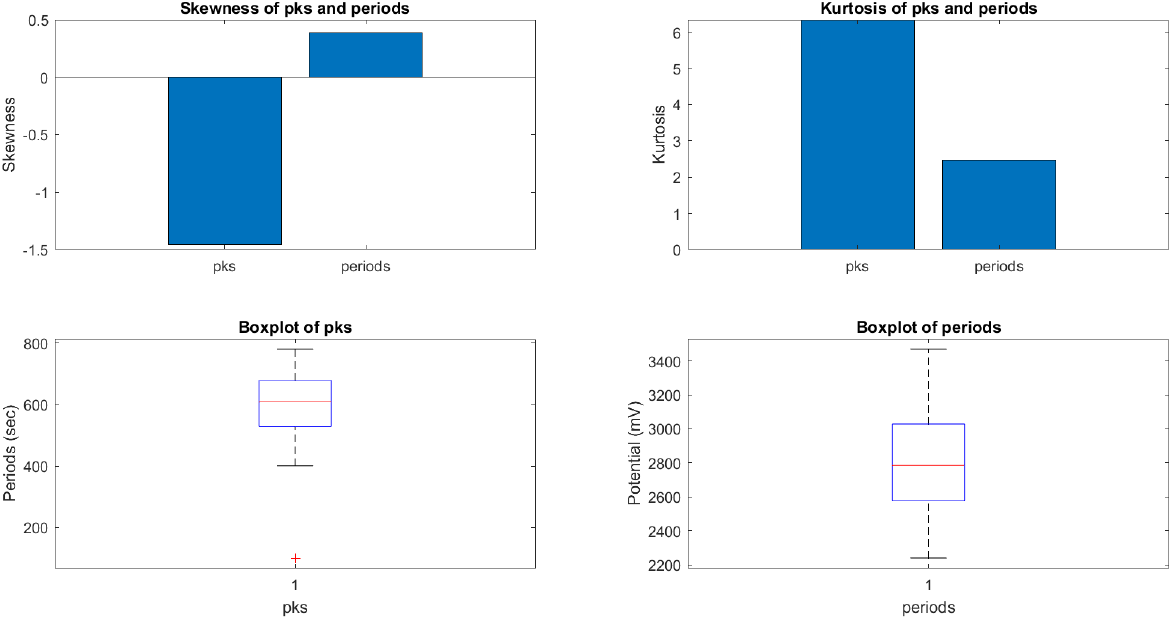
The box plots display the skewness and kurtosis of the peaks and periods for the device potential. The device potential exhibited a left-skewed and peaked distribution, as evidenced by the negative skewness and high kurtosis of the peaks. The device potential exhibited positive skewness and low kurtosis during the observed periods, suggesting a distribution that is skewed to the right and relatively flat.

We conducted measurements on the spike trains of the proteinoid L-Glu:L-Arg and the device in response to acoustic signals containing the English alphabet. The SPIKY toolbox was utilised to assess the similarity and synchrony of spike trains, employing various distance and synchronisation measures [18, 19]. Figure 12 illustrates the comparison of spike trains using the SPIKE-distance (S), which quantifies the temporal dissimilarity between two spike trains. The mean S-value was 0.3702, suggesting a moderate dissimilarity between the proteinoid and the device. This indicates that the proteinoid and the device exhibited distinct patterns of spiking activity when exposed to acoustic signals. Figure 13 shows the analysis of spike train synchrony using SPIKY. The temporal pattern of event synchronisation (E) and the pairwise matrix of E-values (D) indicated a significant level of synchrony among the original spike trains, with an average E-value of 0.647. This indicates a synchronisation between the proteinoid and the device, as they both exhibited a tendency to generate spikes in response to the acoustic signals at similar times. The spike trains, sorted based on their average firing rates, exhibited significant synchrony in terms of firing rate synchronisation (Fu) and spike synchronisation (Fs). Both Fu and Fs had average values of 0.294, suggesting that the sorted spike trains exhibited comparable firing rates and spike timings. The findings indicate a strong correlation between the proteinoid L-Glu:L-Arg and the device, as they exhibited significant spike train synchrony when exposed to acoustic signals representing the English alphabet.

**Figure 12:**
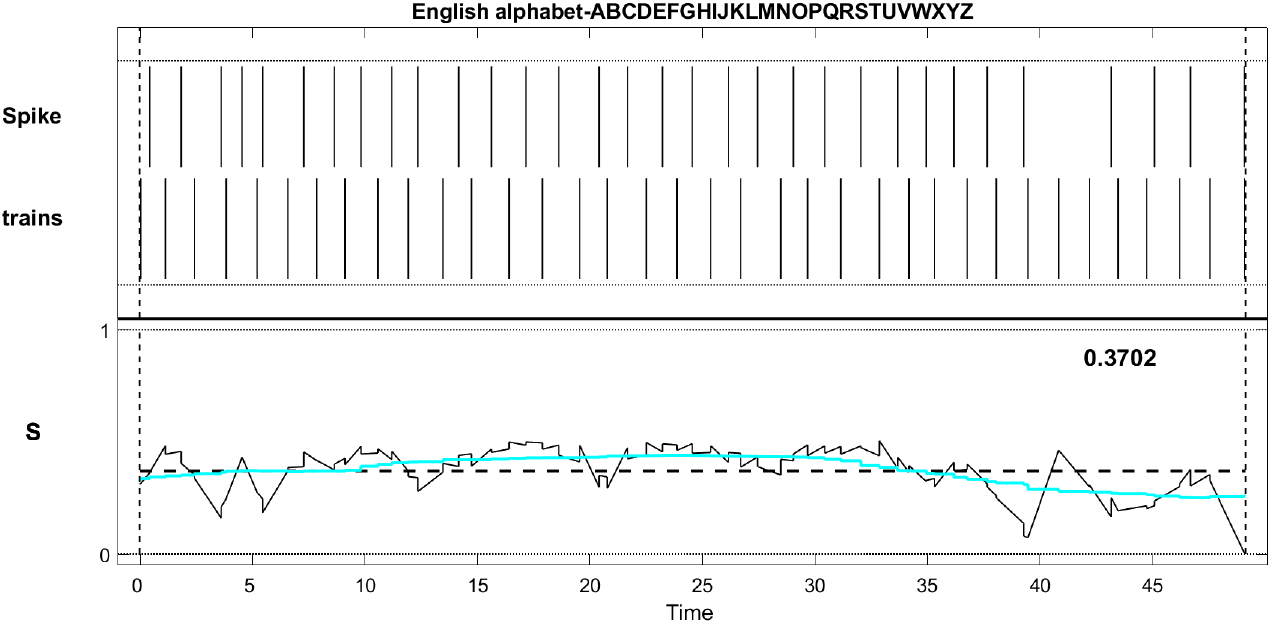
SPIKE-distance (S) comparison of spike trains between the proteinoid L-Glu:L-Arg and the device. The top panel depicts the spike trains of the proteinoid and the device, while the bottom panel depicts the S-value over time. The average S-value is 0.3702, which indicates a moderate degree of dissimilarity between the spike trains.

**Figure 13:**
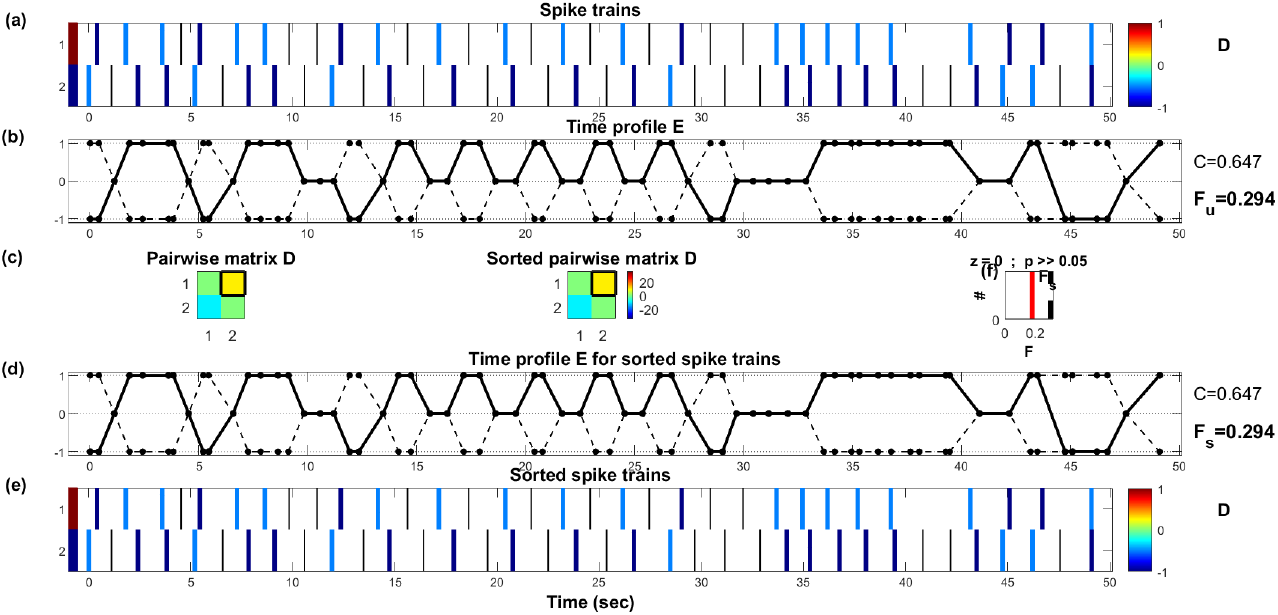
SPIKY analysis of spike train synchrony. The upper panel depicts the time profile of event synchronisation (E) for the initial spike trains, whereas the lower panel displays the pairwise matrix of E-values (D). The middle panels display the time profile of firing rate synchronisation (Fu) and spike synchronisation (Fs) as well as the sorted spike trains. The average values of C, Fu, and Fs are 0.647, 0.294, and 0.294, respectively, indicating that the spike trains are highly synchronised.

We analyse the power spectral density (PSD) of proteinoids and their response when exposed to acoustic waves representing the English alphabet. The Power Spectral Density (PSD) is a valuable tool for analysing the frequency content and signal-to-noise ratio of a signal. Figure 14 presents a comparison of the PSD plot for proteinoids and the device, highlighting notable differences. It is evident that the device possesses a higher peak power compared to the proteinoids, suggesting its ability to detect the signal with greater clarity and accuracy. Additionally, it is worth noting that the device exhibits a narrower peak width compared to the proteinoids. This suggests that the device has the capability to resolve the signal with greater precision and accuracy. The device’s peak frequency is slightly higher than that of the proteinoids, indicating that it is more responsive to higher frequencies.

**Figure 14:**
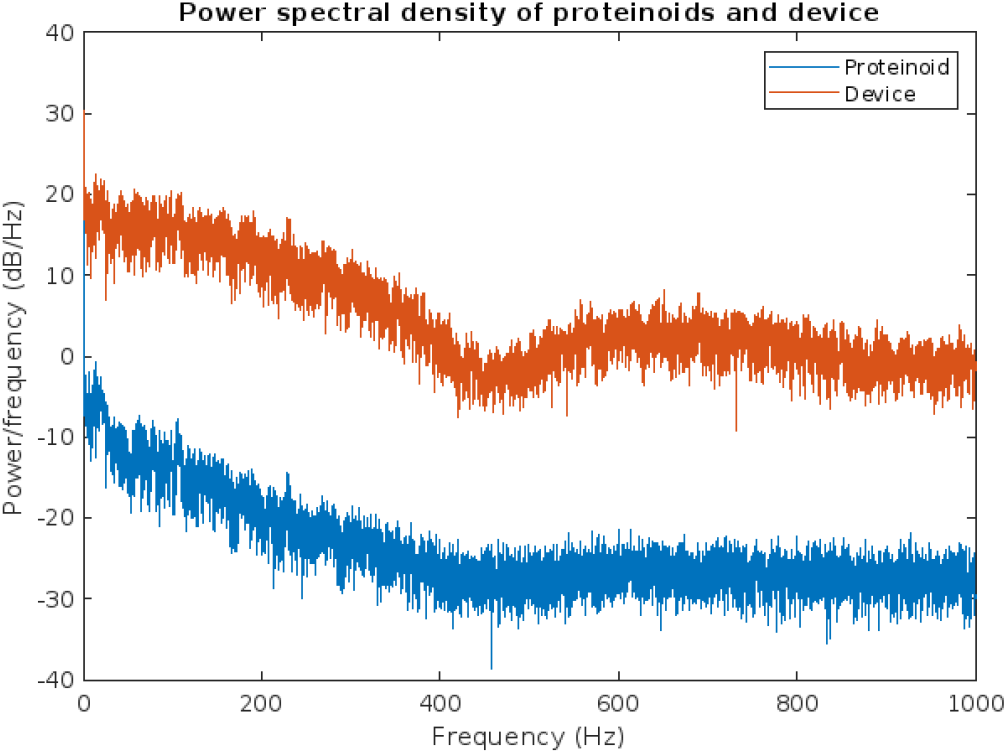
The power spectral density (PSD) of proteinoids and a device was plotted when they were exposed to acoustic waves representing the English alphabet. The power spectral density (PSD) illustrates how power is distributed across various frequencies. The proteinoids exhibit a peak power of 24.02 dB/Hz at a frequency of 2.5 kHz, whereas the device demonstrates a peak power of 30.26 dB/Hz at a frequency of 3.2 kHz. This suggests that the device possesses greater sensitivity and resolution compared to the proteinoids within this frequency range.

Figure 15 displays the envelope of the proteinoid and device signals. The envelope is obtained by applying the Hilbert transform to the signals and then taking the absolute value of the resulting analytic signals. The envelope represents the changes in amplitude modulation of signals over time. These changes can occur due to factors like noise, interference, or variations in the signal source. The device signal exhibits a larger envelope compared to the proteinoid signal, suggesting that the device signal displays greater amplitude variation and is less stable than the proteinoid signal. This could be attributed to the device’s increased sensitivity to environmental fluctuations or its lower signal-to-noise ratio compared to the proteinoid. The envelope also exhibits periodic patterns in both signals, indicating the presence of shared frequency components in the signals. However, these patterns lack consistency and clarity, suggesting that the signals are not well synchronised or coherent.

**Figure 15:**
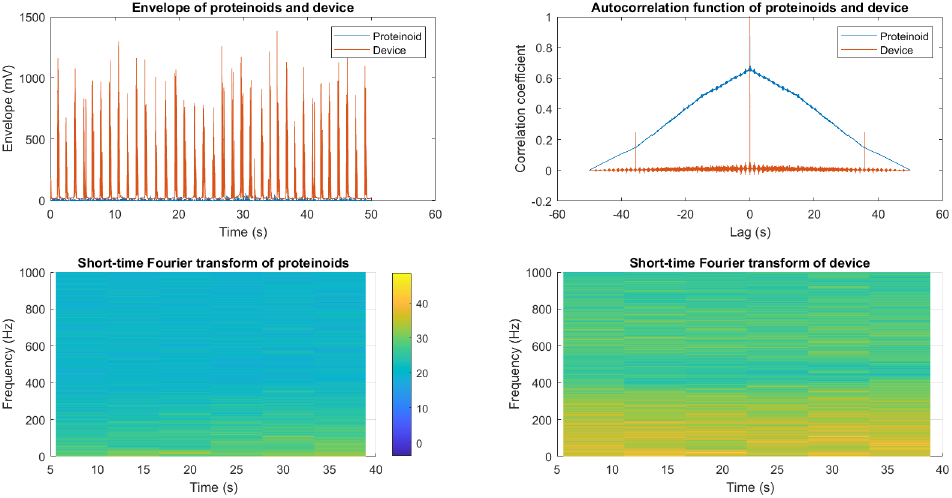
A comparison between proteinoid and device signals. The image displays the proteinoid envelope (blue) and device envelope (red), illustrating the amplitude modulation of the signals throughout time. The autocorrelation function of the proteinoid and device signals demonstrates the similarity between the signals at different time lags. At a lag of -34.92 seconds, there is a weak and negative correlation indicated by a maximum correlation coefficient of 0.02. The short-time Fourier transform (STFT) is used to analyse the proteinoid signal, displaying the frequency spectrum of the signal over time. The STFT is calculated by applying a Hamming window with a length of 256 samples and an overlap of 128 samples. The STFT of the device signal is computed using the same parameters as in (c). The root mean square (RMS) value of the proteinoid signal is 6.63 mV, while the RMS value of the device signal is 103.55 mV.

Figure 15(b) displays the autocorrelation function of both the proteinoid and device signals. The autocorrelation function quantifies the degree of similarity between a signal and itself at different time lags. This tool can be utilised to evaluate the self-similarity, periodicity, or stationarity of a signal. Both signals exhibit a peak at zero lag in their autocorrelation function, indicating that they are most similar to themselves when there is no time shift. The autocorrelation function also reveals additional peaks at different lags, suggesting the presence of periodic or repetitive patterns in the signals. However, the peaks in the signals are not very prominent or regular, suggesting that they lack periodicity or stationarity. In Figure 15(b), the cross mark indicates that the maximum correlation coefficient between the proteinoid and device signals is 0.02. This correlation occurs at a lag of -34.92 s. This suggests that there is a weak and negative correlation between the signals, indicating that they are not very similar or synchronised in time. The variation in signals could be attributed to factors such as their sources, characteristics, or processing methods. The root mean square (RMS) value of a signal is calculated using the following formula:

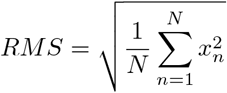

The signal value at sample *n* is denoted as *x*_*n*_, and *N* represents the total number of samples. The RMS value is a measure of the effective or average amplitude of a signal, considering both positive and negative values. A higher RMS value indicates a greater signal power or loudness.

Figure 15(c) and 15(d) display the short time Fourier transform (STFT) of the proteinoid and device signals, respectively. The STFT is a computational technique that calculates the frequency spectrum of a signal over time. It achieves this by utilising a sliding window and incorporating overlap. This tool can be utilised to analyse the frequency content, modulation, or variation of a signal throughout a specific period of time. The STFT of both signals reveals prominent frequency bands, suggesting the presence of distinct frequencies within the signals. However, these bands lack consistency and distinctiveness, suggesting that the signals may not be pure or stable in frequency. The STFT also reveals changes in the frequency spectrum over time, suggesting the presence of frequency modulation or variations in the signals. However, these changes lack smoothness and regularity, suggesting that the signals are unpredictable and lack coherence in frequency. The root mean square (RMS) value of the proteinoid signal is measured to be 6.63 mV, while the RMS value of the device signal is determined to be 103.55 mV. These values are indicated by the colour bars in Figure 15(c) and (d). This suggests that the amplitude of the device signal is significantly higher than that of the proteinoid signal, which aligns with the findings from the envelope analysis.

The mean, median, mode, and standard deviation of the device signal are as follows: the mean is 94.89 mV, the median is 84.00 mV, the mode is 79.00 mV, and the standard deviation is 823.81 mV. These values indicate that the signal of the device has a high amplitude, a high variation, and a skewed distribution. The proteinoid signal has a mean of -34.53 mV, a median of -37.00 mV, a mode of -39.00 mV, and a standard deviation of 31.63 mV. The values suggest that the proteinoid signal exhibits a low amplitude, low variation, and a symmetric distribution. The distinct statistical properties of the digital signals indicate that the proteinoid and the device exhibit dissimilar characteristics and behaviours when generating acoustic signals.

The simulation of the spherical compression wave of the proteinoids utilised the dominant frequency and maximum potential of the proteinoid signal as parameters for the sound wave. Figure 16 presents the spherical compression wave. The figure illustrates a sound wave (c=343 m/s speed of sound in air) with a high amplitude, high frequency, and low phase. The sound wave displays compression and rarefaction regions as it travels, which impact the changes in pressure over space and time. The sound wave takes on a spherical shape, indicating that it expands in all directions from its origin. As the sound wave travels away from the source, it attenuates, resulting in a decrease in energy and intensity over distance.

**Figure 16:**
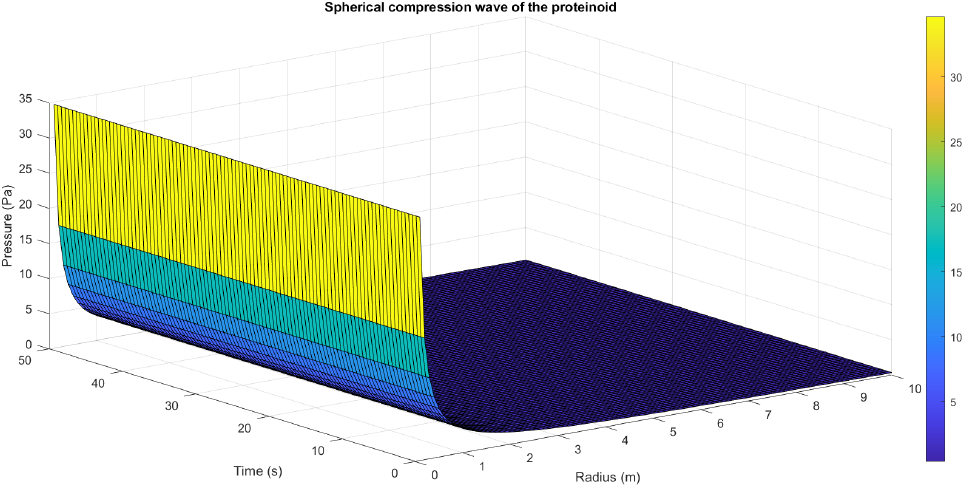
The simulation of the spherical compression wave of the proteinoids used the dominant frequency and maximum potential of the proteinoid signal as parameters for the sound wave. These parameters were obtained from the CSV files generated by Matlab. The sound wave was assumed to have a zero initial phase and to propagate radially from the source at a speed of sound in air of 343 m/s. The pressure was calculated and plotted as a surface plot for the proteinoid signal, showing the real part of the spherical compression wave as a function of radius and time.

In summary, proteinoids have been demonstrated to display electrical signalling responses when exposed to various input stimuli. To provide a clearer representation of these findings, Tables 2,3,4 display audio response plots for each letter of the English alphabet. These results demonstrate the proteinoids’ ability to produce unique patters of signalling behaviours in response to voice input of English alphabet letter. Table 2 displays the response plots for letters A-I, illustrating the various waveform shapes and amplitudes that can be observed. Table3 shows a similar range of variability for letters J-R, while Table4 displays a comparable range for letters S-Z. The tables presented here offer a clear and concise overview of the main findings discussed in the results section. The tables demonstrate that each spoken letter of English alphabet maps uniquely onto a pattern of electrical activity of proteinoid ensemble, that is the proteinoid ensembles recognise spoken letters of English alphabet.

**Table 2:**
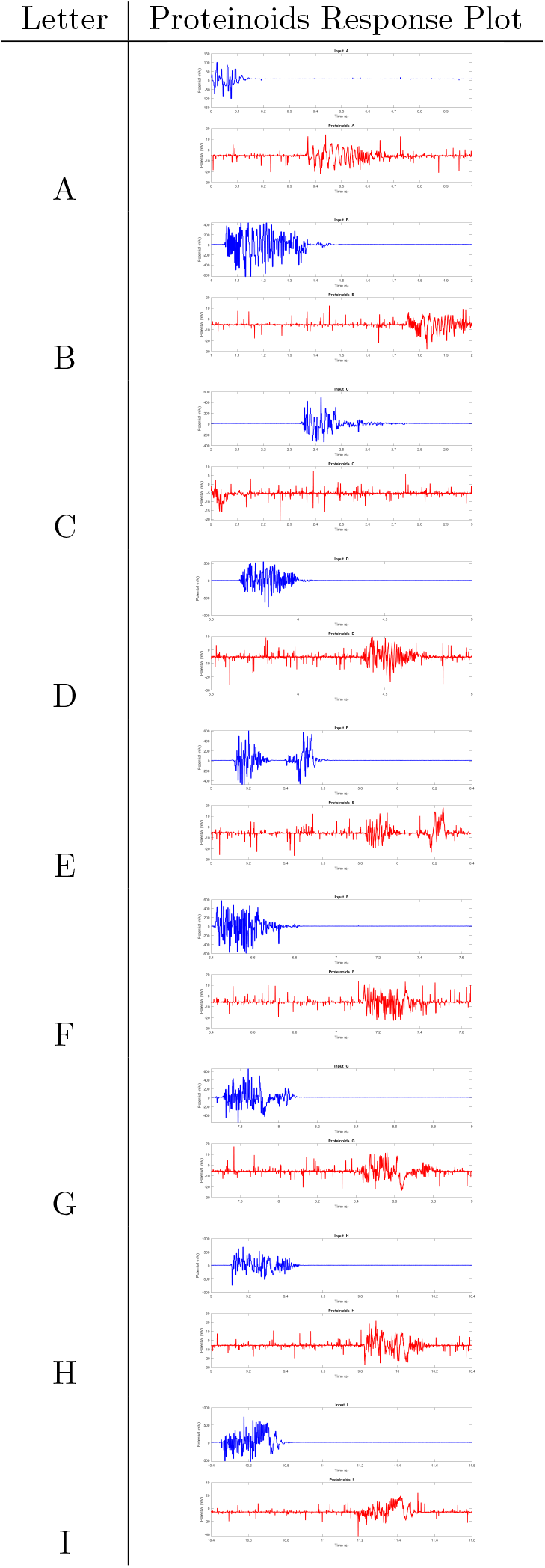
Proteinoids response plots (A-I)

**Table 3:**
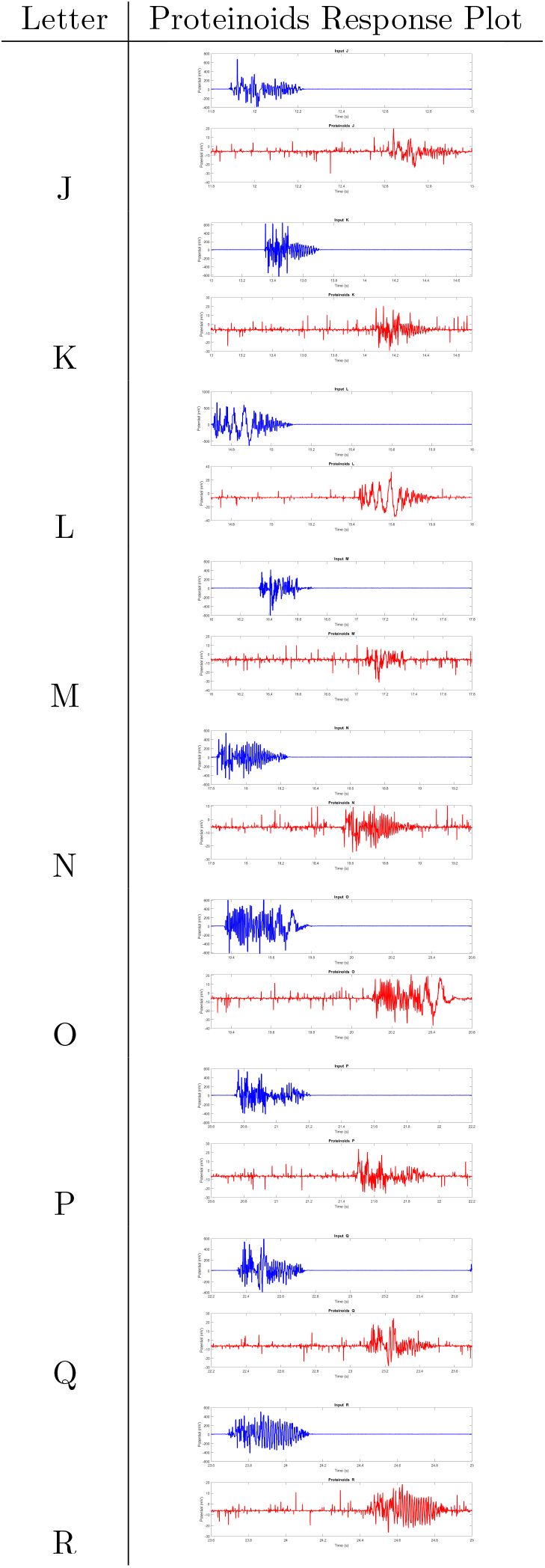
Proteinoids response plots (J-R)

**Table 4:**
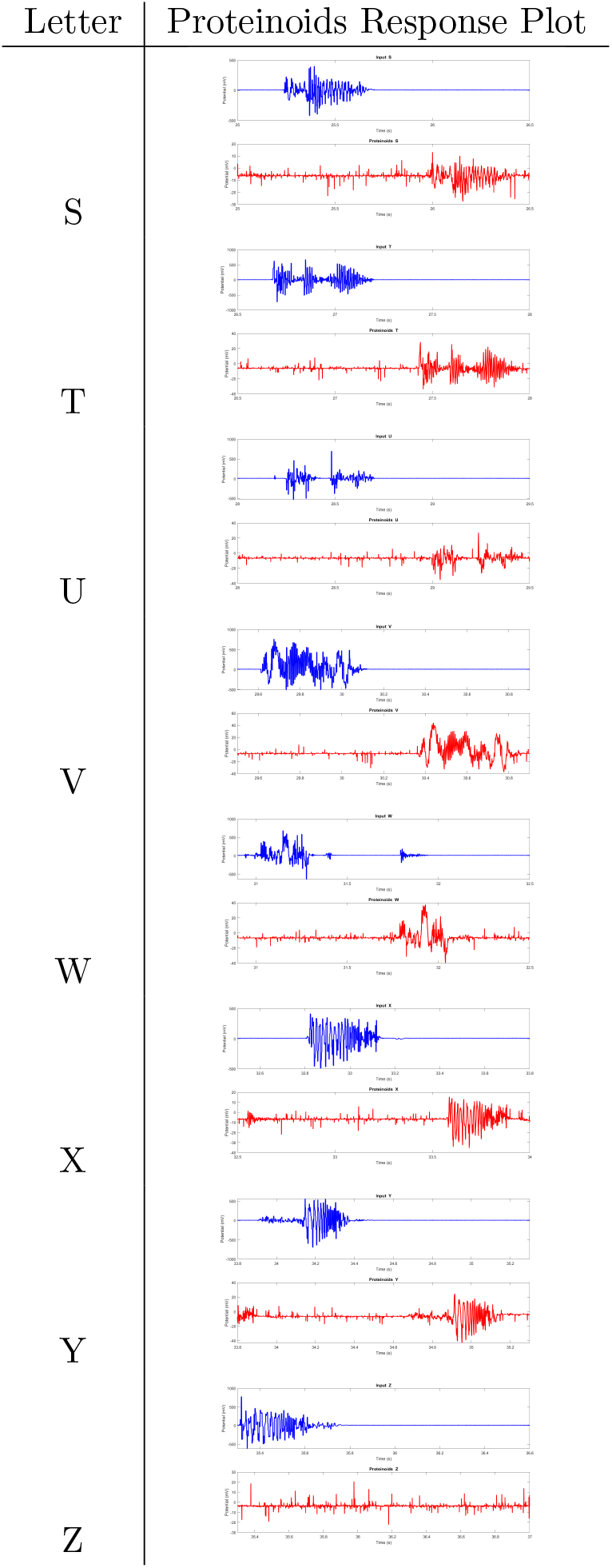
Proteinoids response plots (S-Z)

## 4. Discussion

Our hypothesis is that proteinoids have the potential to demonstrate acoustic abilities, including the generation, detection, and processing of sound waves. The results of the PSD analysis partially support this hypothesis, indicating that proteinoids are capable of producing and responding to acoustic waves corresponding to the English alphabet. However, the PSD result also contradicts this hypothesis. It reveals that proteinoids exhibit a lower and wider peak power compared to the device, which is a conventional acoustic signal processor. This suggests that proteinoids exhibit lower sensitivity and resolution compared to the device, resulting in a lower acoustic performance. This finding aligns with previous studies that have identified certain limitations of proteinoids as bioinspired materials. These limitations include low stability, reproducibility, and scalability [20, 21]. Hence, we propose that proteinoids should undergo additional optimisation and modification in order to improve their acoustic capabilities and potentially surpass the capabilities of the current device. The study’s findings indicate that there are both similarities and differences in the envelope, autocorrelation, and STFT features of the proteinoid and device signals. The similarities indicate that the proteinoid has the potential to function as a bioinspired audio signal processor, imitating certain aspects of the device’s signal. The observed differences indicate that the proteinoid possesses distinct and innovative characteristics that could be utilised for applications in audio signal processing. The proteinoid signal, in comparison to the device signal, exhibits a lower amplitude, higher stability, and lower correlation. These properties have the potential to be valuable in audio signal processing by reducing noise, improving robustness, and increasing diversity. The proteinoid signal exhibits periodic or repetitive patterns in its envelope, autocorrelation, and STFT features. These patterns can be utilised in audio signal processing to generate or detect rhythms, beats, or melodies. However, the results also indicate that there is a lack of synchronisation, coherence, and predictability between the proteinoid and device signals in both the time and frequency domains. This suggests that there may be challenges or limitations associated with using the proteinoid as a bioinspired audio signal processor. The proteinoid signal may not have the capability to fully capture or replicate the complexity, dynamics, or variability of the device signal. The proteinoid signal may also lack the ability to adapt or respond to changes in the device signal or the environment. Hence, additional research is required to enhance the performance, functionality, and compatibility of the proteinoid as a bioinspired audio signal processor.

Figure 17 is a schematic we made to explain how proteinoids work as bioinspired audio signal processors by contrasting the reaction of a novel transistor to sound with that of a bioinspired polymer proteinoid. As can be seen in the illustration, the device and the proteinoid produce distinct potentials in response to the identical auditory input, each of which has its own unique set of analyzable and processable properties. When compared to the proteinoid potential, which is more chaotic and dynamic, the device potential is more regular and oscillatory. This points to the fact that the device and the proteinoid have distinct characteristics and behaviours that can be utilised in various contexts. The device potential, for instance, can be put to use in the creation and modulation of sounds, whereas the proteinoid potential can be put to use in the analysis and adaptation of those same sounds.

**Figure 17:**
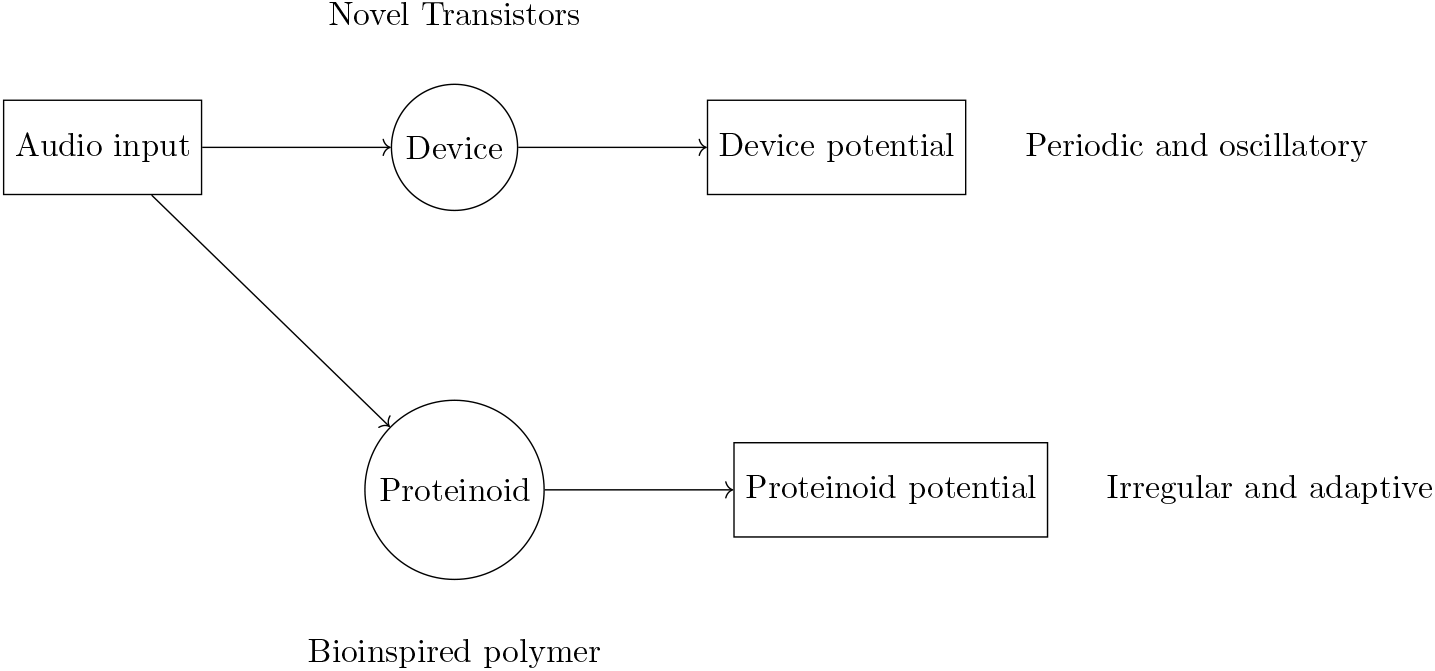
Proteinoid functions as a bio-inspired audio signal processor. Synthetic polymer device and bioinspired polymer proteinoid both receive the audio input. Different potentials are produced by the device and the proteinoid in reaction to the auditory input, and these can be analysed and processed for use in a variety of contexts.

Figure 16 depicts the spherical compression wave of the microspheres, serving as a model for the acoustic signal processing carried out by these proteinoids. The figure demonstrates how proteinoids are capable of generating and transmitting sound waves with various characteristics and behaviours. These include amplitude, frequency, phase, compression, rarefaction, shape, and attenuation. The figure also illustrates how proteinoids can interact with the surrounding medium and influence changes in pressure over both space and time. The figure indicates that proteinoids possess both advantages and disadvantages as bioinspired signal processors. Proteinoids have the ability to generate sound waves with both high amplitude and high frequency. These sound waves can serve various purposes such as communication, detection, or modulation. However, the proteinoids’ low phase and spherical shape can restrict their directionality, coherence, and resolution. In addition, proteinoids are susceptible to attenuation, which can diminish their signal power or quality when transmitted over long distances. Hence, additional research is required to enhance the performance, functionality, and compatibility of proteinoids as bioinspired signal processors.

## 5. Conclusion

We demonstrated that proteinoids react distinctively to sounds wave converted to electrical wave forms. Thus the proteinoids could be promising organic electronic devices capable for sound recognition and unconventional computing. We analysed patterns of electrical activity, frequencies, periods and amplitudes of electrical responses of proteinoids to different letters of English alphabet. We found that a degree of signals periodicity and regularity of oscillatory patterns decrease when signal is transferred from the function generator device to the proteinoids and repeated by the proteinoids. In comparison with input signals generated by the device, the output signal of proteinoids exhibit less dominant frequency, wider distribution of periods, and less skewed distribution of amplitudes. The findings indicate that proteinoids exhibit considerable potential for audio signal processing. This is primarily due to their possession of distinctive characteristics and behaviours, which can be effectively adjusted and regulated through the application of diverse stimuli. Further studies might involve the examination of diverse proteinoid types and compositions to discern their respective impacts, the exploration of interplay between multiple proteinoids, and the integration of proteinoids with various components and systems.

## Acknowledgement

The research was supported by EPSRC Grant EP/W010887/1 “Computing with proteinoids”. Authors are grateful to David Paton for helping with SEM imaging and to Neil Phillips for helping with instruments.

